# Mathematically Modeling Inflammation as a Promoter of Tumour Growth

**DOI:** 10.1101/2020.03.08.982918

**Authors:** Kathleen P. Wilkie, Farjana Aktar

## Abstract

Inflammation is now known to play a significant role in tumour growth and progression. It is also difficult to adequately quantify systemic inflammation and the resulting localized effects in cancer. Here we use experimental data to infer the possible contributions of inflammation in a mouse model of cancer. The model is validated by predicting tumour growth under anti-inflammatory treatments, and combination cancer therapies are explored. We then extend the model to consider simultaneous tumour implants at two distinct sites, which experimentally was shown to result in one large and one small tumour. We use this model to examine the role inflammation may play in the growth rate separation. Finally, we use this predictive two-tumour model to explore implications of inflammation on metastases, surgical removal of the primary, and adjuvant anti-inflammatory treatments. This work suggests that improved tumour control can be obtained by targeting both the cancer and host, through anti-inflammatory treatments, including reduced metastatic burden post-surgical removal of primary tumours.

## 1 Introduction

Inflammation is now recognized to promote cancer initiation, promotion, progression, and metastasis, Coussens and Werb (2002), and has been named as a hallmark of cancer, Hanahan and Weinberg (2011). Perhaps the most striking evidence of the power inflammation has to promote cancer is the ability of anti-inflammatory drugs to lower cancer risk and reduce mortality rates for certain types of cancer, including colon and lung, Rayburn et al. (2009). Inflammation induces angiognesis and promotes cell proliferation, stimulating the cancer to grow. It is a systemic response of the body where pro-inflammatory immune cells, such as neutrophils, dendritic cells, and macrophages, preferentially accumulate to localized sites through the CC- and CXC-chemokines and cytokines such as interleukin-1, tumour necrosis factor *α*, and inteferon *γ*. Coussens and Werb (2002) state that inflammation within tumours has become dysregulated and should be “normalized” to decrease tumour-promoting actions, and increase tumour-inhibiting actions of infiltrating immune cells.

Cancers induce chronic inflammation as the pro-inflammatory signals never resolve, resulting in ongoing immune recruitment and increased inflammation, Coussens and Werb (2002). Through alterations to gene expression, enzymes, and transcription factors, chronic inflammation results in multiple pro-cancerous modifications including: increased DNA damage, increased DNA synthesis, cell proliferation, disrupted DNA repair, inhibition of apoptosis, promotion of angiogenesis, and promotion of invasion, Hofseth and Ying (2006). Tumour necrosis further drives the chronic inflammatory environment as necrotic cells produce danger signals (compared to the normal clearance signals produced by apoptotic cells) – and danger signals stimulate a pro-inflammatory state, Chen et al. (2014). Local presence of chemokines and cytokines draw inflammatory cells out of the blood stream through chemotaxis. Both immune and cancer cells produce these recruitment signals, potentially creating a stronger chemotactic draw at larger tumours than at smaller ones.

Measurments of systemic inflammation include cell counts of neutrophils or lymphocytes, blood levels of acute phase proteins such as C-reative protein; or cell ratios such as the neutrophil lymphocyte ratio, Roxburgh and McMillan (2014). Such measurements, however, have inconsistent reports on their prognostic value and are easily confounded by comorbidities, Sylman et al. (2018). Nevertheless, inflammation has become a target for both cancer prevention and therapy, with cyclooxygenase 2 (COX-2 / PTGS2) being the most common anti-inflammatory target, Rayburn et al. (2009).

In this work, we explore the role of inflammation in tumour growth, through a single and a simultaneous double tumour experiment from Benzekry et al. (2017). Their experimental data of Lewis Lung Carcinoma (LLC) tumours in the C57BL6 mouse, show that with two simultaneous tumours, one consistently grows larger than the other, even though the experimental implants are the same. They proposed concomitant resistance as a possible explanation for these observations. Here, we propose an alternate hypothesis based on the idea of unbalanced accumulation of tumour-promoting inflammation, and use our mathematical model to study the role of inflammation in both the single and double tumour experiments. Indeed, Williams et al. (2000) showed that the anti-inflammatory (COX-2 inhibitor) Celecoxib inhibits LLC tumours in C57BL6 mice, demonstrating that inflammation plays a role in this mouse model.

The phenomenon of concomitant resistance (CR) describes the appearance of secondary tumours to have delayed growth dynamics compared to the primary. This has been applied to the study of metastases, and the observation that metastases can experience significant increases in growth following primary tumour resection, Chiarella et al. (2012). Three main hypotheses have been proposed to explain CR. The first theory is concomitant immunity, which proposes that the primary tumour generates an anti-tumour response sufficient to inhibit smaller secondary tumours while not effectively inhibiting the primary, Bashford et al. (1908). This hypothesis does not explain CR observed in weak to non-immunogenic tumours, Ruggiero et al. (1990); Chiarella et al. (2012); Gorelik (1983), such as the LLC–C57BL6 model system, Lechner et al. (2013), used in the Benzekry et al. (2017) experiments. The second theory proposes that CR may result from nutrient competition, in that an established tumour traps glucose, nitrogen and other nutrients, Tyzzer (1913); Gorelik (1983), leaving little to support the secondary tumours. Lastly, the third hypothesis suggests that the primary tumour may produce anti-angiogenic or anti-proliferative substances that suppress the secondary implant, Ruggiero et al. (1990); O’Reilly et al. (1994); Gorelik (1983). Indeed O’Reilly et al. (1994) demonstrated that murine LLC tumours could inhibit metastases through production of angiostatin. And it has recently been found that meta- and ortho-tyrosines can inhibit cancer cell proliferation while large tumour accumulation of certain amino acids counteracts these inhibitory effects, creating the observed CR, Ruggiero et al. (2011); Chiarella et al. (2012). Together, these last two hypotheses can explain CR in non-immunogenic tumours but not the specific inhibition of secondary implants observed in immunogenic tumours, Chiarella et al. (2012).

Additionally, mechanisms of CR appear to work on different timescales. Early CR (small tumours) may be attributed to tumour immunogenicity, mid CR to limited angiogenesis, and late CR (large tumours) to inhibited cell proliferation, Chiarella et al. (2012); O’Reilly et al. (1994). Since the experimental set-up referenced here involves simultaneously injected (and nominally equal in size) non-immunogenic tumours, none of the CR mechanisms seem to directly apply.

Hence, we propose that inflammation may play a role in the observed growth differences. In simultanteously growing tumours at distant sites, initially small discrepancies in their size, inflammatory state, or microenvironment may grow into significant differences. This may be driven by preferential accumulation of inflammatory cells to one tumour site over the other, via positive feedback loops in signaling networks and signal amplification from cancer cell production of pro-inflammatory cytokines. For example, unbalanced immune chemoattractors (such as CCL19/CCR7, CXCL9/CXCR3, IL15/IL15R) has been observed between breast metastases and the primary tumour, Szekely et al. (2018).

In breast cancer, the relapse risk following surgery is bimodal, with the first peak generally attributed to surgery-induced inflammatory acceleration of metastases from a dormant state (with removal of CR mechanisms), and the second peak generally attributed to stochastic transition of metastases from the dormant to proliferative state, Galmarini et al. (2014). Indeed, the first peak can be suppressed by administration of a non-steroidal anti-inflammatory drug (NSAID) in the perioperative period, Retsky et al. (2012).

Benzekry et al. (2017) proposed that the different growth rates observed in simultaneously injected tumours may be due to CR. They developed mathematical models to explore the hypotheses of limited angiogenesis and inhibited proliferation, and found that the inhibited proliferation model best fit their experimental data. Others models, Poleszczuk et al. (2016); Walker et al. (2018) explored growth disparities under the hypotheses of immune cytotoxicity and blood flow fractions. These models set different blood flow fractions to different sites, and see that increased blood flow can lead to disparate growth rates due to immune cytotoxicity. Here, we consider tumours growing in (assumed identical) regions - both sides of the caudal half of the back, so blood flow is nominally equal. Also, instead of immune cytotoxicity, which is limited in the LLC–C57BL6 model, Lechner et al. (2013), we consider immune stimulation through inflammation.

To explore the role of inflammation in cancer, we present a mathematical model of tumour growth in a pro-inflammatory state. Using ordinary differential equations, our model describes the growth of each compartment: cancer, cancer carrying capacity, and immune volumes. We choose this type of model as it is a common and particularly useful method to describe cancer-immune interactions, Adam and Bellomo (1997); Eftimie et al. (2011); Kuznetsov et al. (1994); de Pillis et al. (2005); Wilkie and Hahnfeldt (2013); Wilkie (2013); Arciero et al. (2010). Most mathematical models of cancer-immune interactions forego direct stimulatory actions of the immune response and only consider the cytotoxic actions. This is primarily because immune predation is a potential mechanism to achieve tumour elimination, and it can be modulated by immunotherapies.

Inflammation, however, is now accepted as a tumour-promoter, and can also be targeted, so is thus a valuable addition to model formulations. In light of this, we have worked to directly incorporate such promotion into mathematical models, Wilkie and Hahnfeldt (2017). The model we propose here allows for inflammation-driven stimulation of tumour growth through increases to the environmental carrying capacity. Other models include immune-driven stimulation by increasing the basal cancer growth rate directly, den Breems and Eftimie (2016); Louzoun et al. (2014).

The paper proceeds as follows. A single tumour model is presented and model parameters are estimated by fitting to experimental data. Stability analysis is discussed, and parameter sensitivity is examined locally. The model is then extended to a two-tumour bearing host model, and similar stability analysis is presented. Next, the significance of inflammation to the LLC–C57BL6 mouse model is explored by comparing model predictions to experimental data in mice with restricted inflammatory responses. Treatment efficacy on a single tumour is then presented, first by combining chemotherapy with anti-inflammatories, and then by including anti-angiogenic drugs. Next, we return to the two-tumour model and explore the role inflammation may play in the growth rate differences observed in experimental data. We then use this two-tumour model to examine the metastatic state and effects of surgery, and possible adjuvant anti-inflammatory therapy, on secondary tumours.

## 2 Mathematical Model and Assumptions

Here we develop a mathematical model to describe inflammation-driven tumour growth in both single and double tumour-bearing hosts.

### 2.1 Experimental Data

Benzekry et al. (2017) reported experimental data that describes murine Lewis Lung Carcinomas (LLC) grown in C57BL6 mice. Subcutaneous injections of 10^6^ LLC cells in 0.2 ml phosphate-buffered saline were injected into anesthetized mice on the caudal half of the back in the control group, and on the two lateral sides of the caudal half of the back in the two-tumour group. Measurements were taken regularly with calipers to a maximum of 1500 mm^3^, when the mice were sacrificed.

Ten mice were included in each group. Consistently, in the two-tumour group, one tumour grew larger than the other, irrespective of left or right sides. Two mice were excluded from our analysis since the injections led to tumours of approximately equal size. These tumours were found to be connected directly by substantial blood vessels upon resection. Since the two tumours were physically connected, they would directly share their inflammatory and other signals and thus significantly reduce any systemic recruitment differences between the two sides. Data is reported as mean ± standard error for all data points, grouping the tumour volumes as either Control, Large, or Small tumour.

### 2.2 Model Equations

The developed model below describes tumour growth with a dynamic carrying capacity in a pro-inflammatory environment. We ignore immune cytotoxicity because the experimental data is from a Lewis Lung Carcinoma (LLC)–C57BL6 mouse model, wherein the cancer grows essentially without any cytotoxic immune activity, Lechner et al. (2013). This is due to the fact that the LLC cell line is derived from a spontanteous lung tumour from a C57BL6 mouse and has thus been immuno-edited to be resistant to immune attack. We do, however, assume that the host has an intact inflammatory response, that is capable of stimulating tumour growth through angiogenesis and production of growth signals.

#### 2.2.1 Single Tumour Model

For one tumour growing in a host, we assume logistic growth with an intrinsic growth rate of *α* and a dynamic carrying capacity *K*_*C*_ (*t*). Where, *C*(*t*) is tumour volume, the rate of growth can be described by:

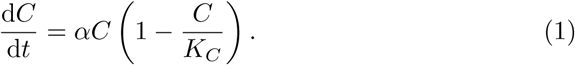

The inflammatory immune response is assumed to grow according to logistic growth as well, with intrinsic growth rate *β* and constant carrying capacity *K*_*I*_. The cancer presence stimulates the inflammatory response, so we include this with an additional term *ρβC*, which is also modified by the carrying capacity limitation. Where *I*(*t*) is immune cell volume at the tumour site, the growth dynamics are thus described by:

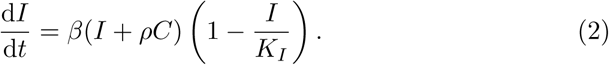

Note that equation (2) models the chronic inflammatory immune response when activated by a growing tumour. The limiting value *K*_*I*_ may be larger for acute immune responses, but here we assume a moderate chronic level of inflammatory action, similar in scope to the tumour. In previous work, Wilkie and Hahnfeldt (2017), a homeostatic maintenance term was added to this equation, to ensure the immune response returned to a healthy level in the absence of a tumour. In this work, however, we neglect the resolving phase of an immune response and assume that the chronic inflammation persists.

While the immune carrying capacity is assumed to be constant, the cancer carrying capacity grows with tumour volume. We follow the formulation derived by Hahnfeldt et al. (1999), but modify it to include inflammatory actions that contribute to both the stimulatory and inhibitory signals controlling environmental capacity, Wilkie and Hahnfeldt (2017). For cancer carrying capacity *K*_*C*_ (*t*) we let *p* be the stimulation coefficient and *q* be the inhibition coefficient. The stimulation term is assumed to be proportional to tumour volume (*V*), while the inhibition term is proportional to the volume raised to the power 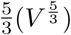, Hahnfeldt et al. (1999). Since *I*(*t*), *C*(*t*), and *K*_*C*_ (*t*) are all measures of volume contributing to the tumour bulk, we describe the cancer carrying capacity growth rate as:

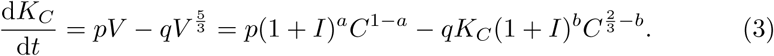

Without loss of generality, the factor (1 + *I*) is used in place of *I* to include a basal level of immune presence and allow for cancer growth without an immune response above this level. Parameters *a* and *b* control the weight placed on immune actions that stimulate and inhibit capacity growth, respectively. Since we are interested in pro-inflammatory actions, we assume *a* > *b* while also restricting 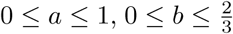, and *b* to be small. To fulfill these requirements, we assume that *a* = 0.2 and *b* = 0.1.

Taken together, equations (1)–(3) describe the growth of one tumour within a microenvironment composed of both cancer and pro-inflammatory immune cells.

#### 2.2.2 Double Tumour Model

To model a host with two simultaneously growing tumours at distant sites, we let *C*_1_ and *C*_2_ denote the two cancer masses, where each volume grows according to equation (1) (with *C* = *C*_*i*_, *i* = 1 or 2) and with dynamic carrying capacities *K*_*C*1_ and *K*_*C*2_ respectively, governed by equation (3) (with *K*_*C*_ = *K*_*Ci*_, *I* = *I*_*i*_, and *C* = *C*_*i*_, *i* = 1 or 2). Since the two tumours are connected through the systemic circulation, they can draw inflammatory cells from the same host reserve. Thus, the immune equation is modified to reflect this competition for inflammatory cells. All together, the equations that describe the two-tumour experiment (with *i* = 1 or 2) are:

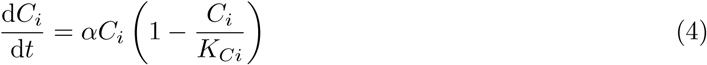

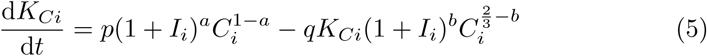

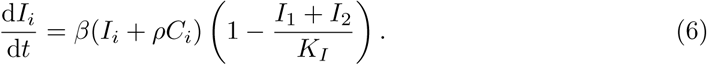

### 2.3 Model Parameterization

Some model parameters were fixed *a priori*, others were fit to experimental data using a simulated annealing algorithm. All parameter values used are listed in Table 1. Initial cancer size was determined by the experimental condition of 10^6^ injected cells which translates to *C*(0) = *C*_0_ ≈ 1 mm^3^. Immune capacity (*K*_*I*_ = 1500 mm^3^) was chosen to approximately match the maximal tumour size in our data. Parameters *a* and *b* are chosen such that a small weight is attributed to immune actions that stimulate tumour capacity (*a* = 0.2) and even less weight is attributed to immune actions that inhibit tumour capacity (*b* = 0.1). Numerical simulations were conducted using Maple and MATLAB software packages.

**Table 1:**
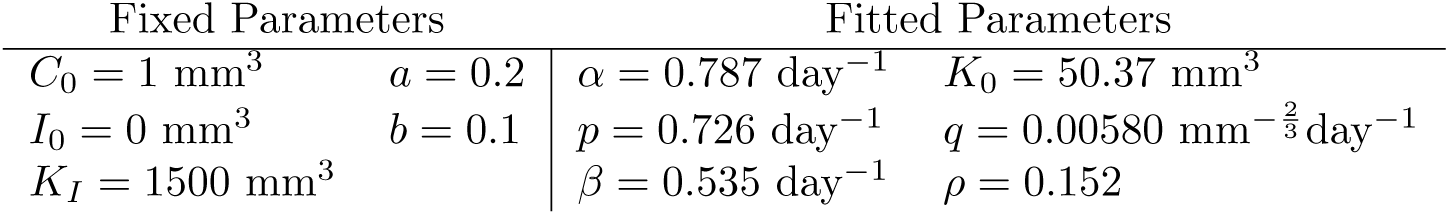
Parameter values for equations (1)–(3). Fixed parameters were set prior to data fitting. Fitted parameters were determined in the simulated annealing algorithm by fitting the model to control experimental data. After ten successful runs, we chose the fit with the smallest goodness-of-fit measure: RMSE = 25.25.

| Fixed Parameters |  | Fitted Parameters |  |
| --- | --- | --- | --- |
| $C_0 = 1 \text{ mm}^3$ | $a = 0.2$ | $\alpha = 0.787 \text{ day}^{-1}$ | $K_0 = 50.37 \text{ mm}^3$ |
| $I_0 = 0 \text{ mm}^3$ | $b = 0.1$ | $p = 0.726 \text{ day}^{-1}$ | $q = 0.00580 \text{ mm}^{-\frac{2}{3}} \text{ day}^{-1}$ |
| $K_I = 1500 \text{ mm}^3$ | | $\beta = 0.535 \text{ day}^{-1}$ | $\rho = 0.152$ |

A simulated annealing algorithm, Goffe et al. (1994); Corana et al. (1987); Marsh et al. (2007), is used to fit the mathematical model, equations (1)–(3), to time-series data of average tumour volume (single tumour bearing control group). The model equations are fit assuming the fixed parameters described in Table 1. This leaves the algorithm to determine the best fit by varying parameters *α, K*_0_, *p, q, β*, and *ρ* by minimizing an objective function, here taken to be the root mean squared error (RMSE). The algorithm is probabilistic and is required to run for at least 15,000 iterations before convergence is accepted, up to a maximum of 35,000 iterations. A brief outline of the algorithm follows:

1. Generate a random initial guess for parameter vector *V* = [*α, K*_0_, *p, q, β, ρ*] and objective function value RMSE.
2. One at a time, perturb each parameter to generate a trial vector *V*′ with a new objective function value RMSE′.
3. Accept this new parameter value if the fit is better (*RMSE*′ < *RMSE*) or based on a probabilistic condition 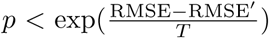 where *T* is the algorithm temperature and *p* is a random number uniformly distributed between 0 and 1.
4. Repeat from 2 until convergence is met. Decrease temperature *T* and refine new guess generations periodically.

## 3 Results

The result of fitting the parameters of the single tumour model, equations (1)–(3), to the control data of Benzekry et al. (2017) is shown in Figure 1(a). Parameter values are listed in Table 1.

**Figure 1:**
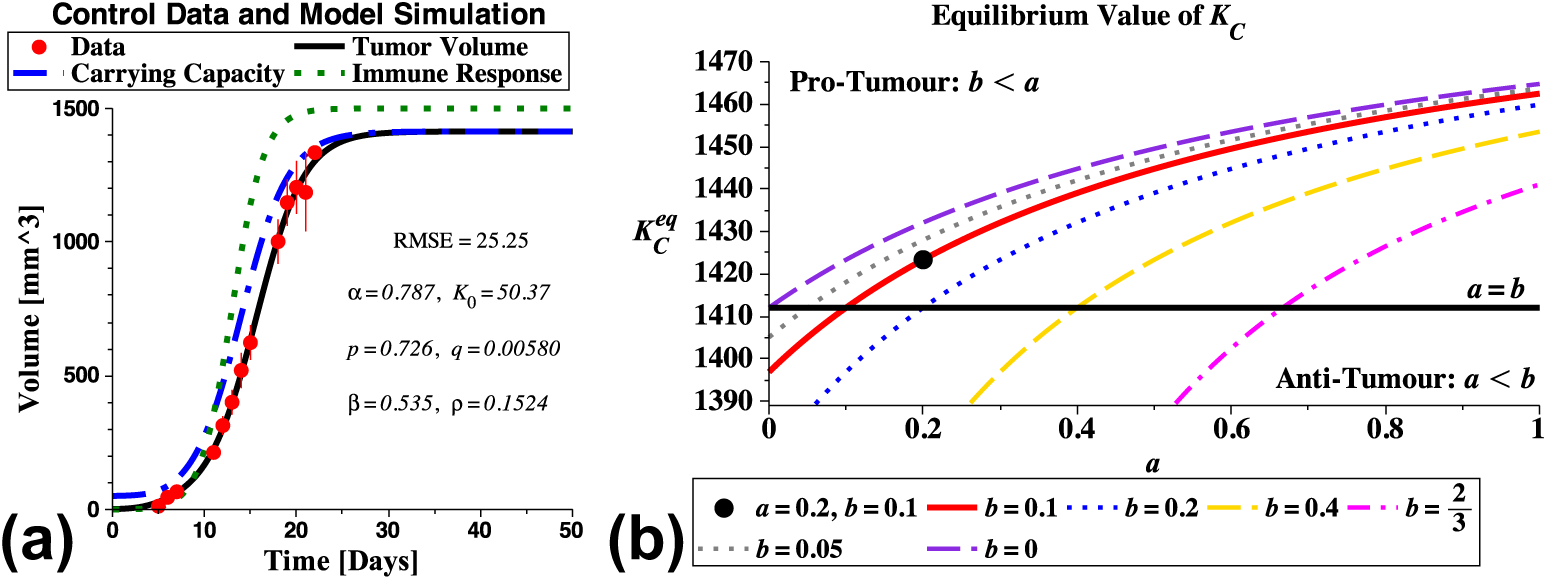
(a) Experimental data (with standard error) for the single tumour-bearing group with model simulation using parameters from Table 1. (b) The dependence of the equilibrium value 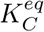 on parameters *a* and *b*, which control the contribution of immune activity to stimulation or inhibition of carrying capacity, respectively. For pro-tumour immune stimulation we require *a* > *b* so that more weight is assigned to the stimulatory immune actions than to inhibitory actions. Here we assume *a* = 0.2 and *b* = 0.1 (indicated by the black dot).

### 3.1 Single Tumour Model Analysis

The mathematical model for one tumour, described by equations (1)–(3) has equilibrium solutions of 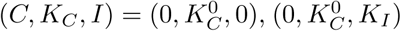, and 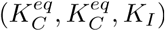. The first two states are unstable cancer-free equilibria with 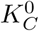 being any initial condition for *K*_*C*_, and the third state is the stable equilibrium representing maximum cancer volume, determined by 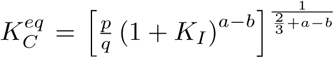. Since the model does not allow cytotoxic immune cell killing of cancer cells, these results are to be expected.

In equation (3), parameter *a* is the weight of immune actions that increase the cancer carrying capacity while *b* is the weight of actions that reduce the capacity. Requiring all exponents to be non-negative implies that 0 ≤ *a* ≤ 1 and 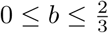. In this work, we assume immune stimulation of cancer growth is the dominating action of immune cells rather than inhibition. This imposes the requirement that *a* > *b*. As these parameters are challenging to estimate from experimental data, we choose to fix them at biologically reasonable estimates. Small values for *a* and *b* ensures that the cancer is the main driving force behind cancer carrying capacity growth. Thus, we choose *a* = 0.2 and *b* = 0.1. The dependence of equilibrium cancer volume, 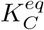, on the values of *a* and *b* is demonstrated in Figure 1(b). The parameter region of interest, where our inequalities on *a* and *b* are satisfied, is the region above the line *a* = *b*. When *b* = 0.1, changing *a* from *a* = *b* to *a* = 1 (the range of acceptable values for *a*) results in an increase in 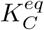 of only 3.6%. The largest change, when *b* = 0, results from varying *a* from 0 to 1, and gives less than a 4% increase in maximum tumour size, demonstrating that the equilibrium behaviour of the model is robust to variations in *a* and *b*.

Model parameter sensitivity is now explored both in terms of predicted tumour volume and tumour growth rate, at day 15. Figure 2 displays model sensitivity plots to the fitted and fixed parameter values listed in Table 1. Figure 2(a) shows that the intrinsic growth rate (*α*) and capacity stimulation coefficient (*p*) most strongly influence tumour volume on day 15, while Figure 2(b) shows that in addition to these two parameters, capacity inhibition coefficient (*q*) most strongly influence the tumour growth rate on day 15. This suggests that tumour dynamics, described by parameter values in Table 1, are strongly regulated by the tumour microenvironment in addition to the cancer’s intrinsic growth rate. Local Sensitivity Coefficients *S*_*i*_, Zi (2011), confirm this result, Figure 2(c). Sensitivity Coefficients are calculated based on the formula

**Figure 2:**
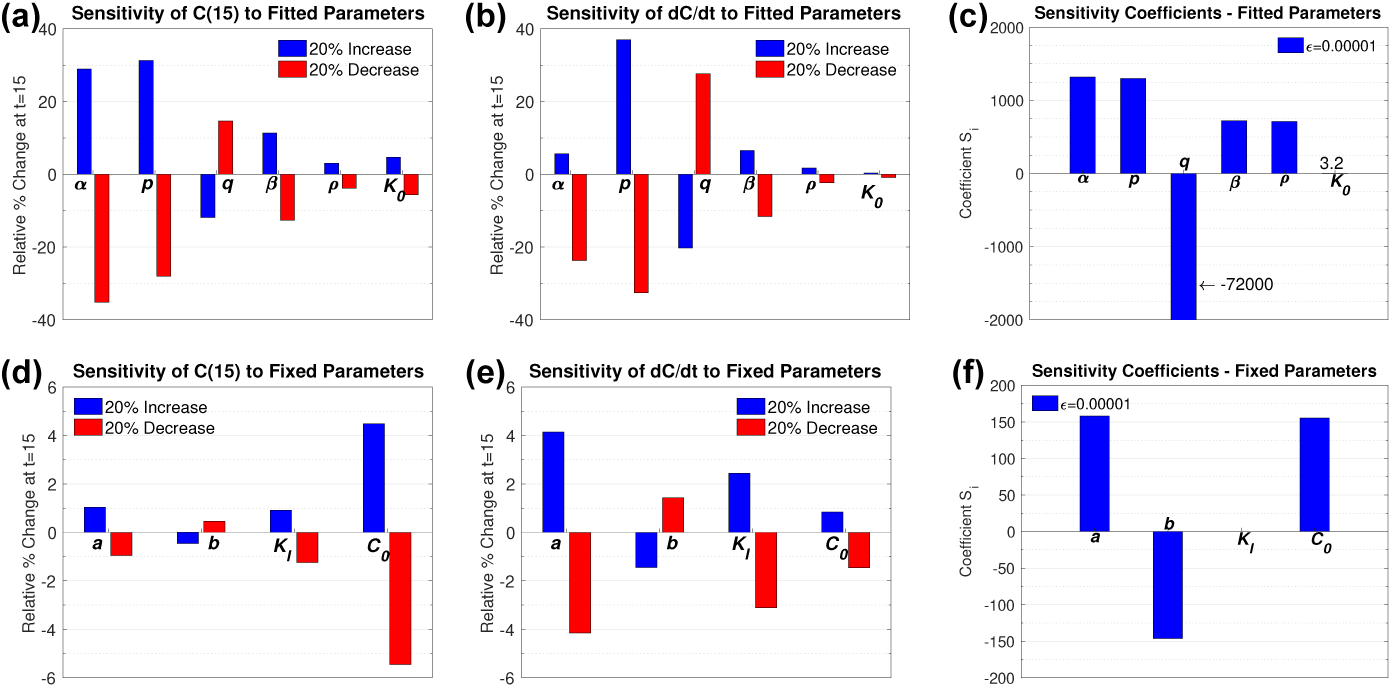
Model sensitivity analysis. Each parameter value is increased or decreased by 20% and relative change in tumour size on day 15 is reported in (a) for fitted parameter values and (d) for fixed parameter values. Similarly, the effect of a 20% increase or decrease on tumour growth rate on day 15 is reported in (b) for fitted parameter values and (e) for fixed parameter values. Sensitivity Coefficients measure local sensitivity of predicted tumour volume to perturbations in fitted (c) and fixed (f) parameter values. Base parameter values are those listed in Table 1.

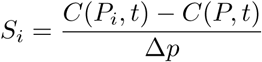

where time *t* = 15 days, 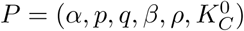 is the vector of parameter values, *P*_*i*_ is this vector with the *i*^*th*^ parameter perturbed by the formula *p* = *p* + Δ*p* = *p* + *p* ∗ *ϵ*.

Figure 2(d-f) show the model sensitivity to fixed parameters (*a, b, K*_*I*_, *C*_0_), and indicate that tumour volume is most sensitive to initial injection size (*C*_0_) while the growth rate on day 15 is most sensitive to parameter *a*, which controls the weight of immune stimulatory actions in increasing tumour carrying capacity. The influence of fixed parameters is generally less than that of fitted parameters, as indicated by smaller sensitivity coefficient magnitudes, suggesting that our choice of parameters to fix was reasonable.

### 3.2 Double Tumour Model Analysis

Plotting the two-tumour experimental data visualizes the growth rate separation that occured in the simultaneously growing tumours, Figure 3(a). One of the tumours is significantly inhibited compared to the control group (small tumour), while the other grows comparable to the control (large tumour).

**Figure 3:**
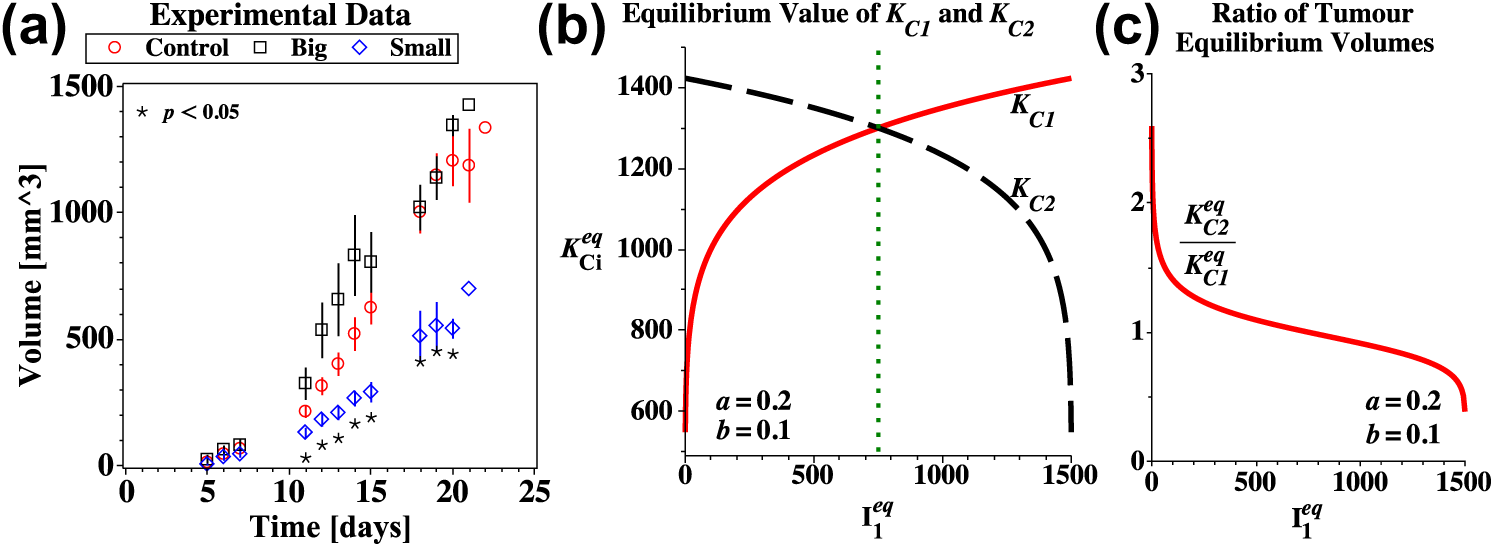
Two-tumour bearing experiment and model steady-state analysis. (a) Average tumour volume for the control and 2-tumour groups, data from Benzekry et al. (2017). Error bars are standard error and statistical significance with *p* < 0.05 is indicated by *. (b) The relation between equilibrium tumour size at each site, as a function of immune presence. One large and one small tumour at equilibrium requires 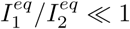 or ≫ 1. (c) The fold-change between the two tumour equilibrium volumes as a function of immune presence at site 1, 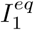.

In our mathematical model, the two simultaneous tumours compete for immune recruitment, equation (6). The cancer and cancer carrying capacity equations, (4) and (5), become specified for either site 1 (*C*_1_(*t*) and *K*_*C*1_(*t*)) or site 2 (*C*_2_(*t*) and *K*_*C*2_(*t*)). And each site draws their own immune component (*I*_1_ or *I*_2_) from the pool (*K*_*I*_). Again, the cancer-free equilibria are unstable states:

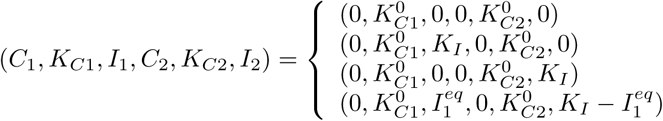

The equilibrium states involving only one of the tumours reaching maximum size are also unstable states:

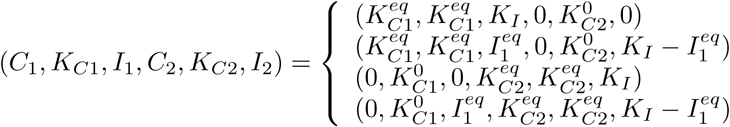

Finally, the only stable state is the equilibrium with two tumours:

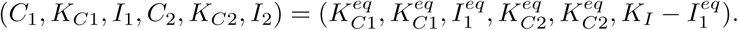

The maximum cancer volume at each site depends on immune presence at that site:

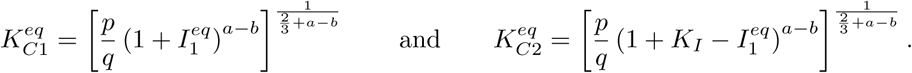

This dependence is demonstrated in Figure 3(b). Figure 3(c) shows the equilibrium tumour volume imbalance as a function of immune presence 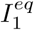. In long time, the tumours achieve approximate equality except in extreme cases of immune imbalance.

Equilibrium solutions represent long-time behaviour and give no information about growth dynamics before then. This model suggests, that even though the growth rates are different between the two sites for early times, their equilibrium sizes will be approximately equal (ratio being about 1) in all but extreme cases, Figure 3(c). Note that in an LLC tumour the equilibrium sizes are likely larger than the host mouse can sustain, and thus are not reached in their lifetime. This is especially true in experiments for ethical reasons, and a consequence is the underestimation of final tumour size in parameterization. For simulation purposes, however, we assume the maximum tumour size about 1500 mm^3^. The extreme case where almost all immune cells localize to one site over the other seems improbable given the experimental set-up of equal injections. In such cases, the model predicts only a 2.5-fold difference in equilibrium tumour volume. In the limiting case, where 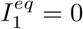 and 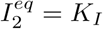, the ratio

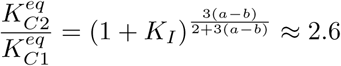

for parameter values listed in Table 1. Thus, while the growth rates are quite different between the two sites for early times, this model suggests that the smaller tumour will ultimately catch up to the larger one in long time (provided the host survives this long).

### 3.3 Inflammation Promotes Tumour Growth

Here we explore the role of inflammation in the LLC-C57BL6 tumour mouse model. The COX-2 enzyme promotes tumour growth through a sequence of events that produce vascular endothelial growth factor (VEGF), and it is a common target of many NSAIDs, Rayburn et al. (2009). Williams et al. (2000) showed that LLC tumour growth in a C57BL6 mouse can be suppressed by administering a selective COX-2 inhibitor (Celecoxib) or by genetically knocking out COX-2, in either a COX-2^+*/*−^ (partial) or a COX-2^−*/*−^ (complete) mouse. Using Williams’ data we can validate our model predictions for the dependence of this tumour/mouse model on inflammatory actions. To do so, we numerically compute solutions to the single tumour model, equations (1)–(3), with the following conditions:

1. Control: parameter values listed in table 1 and *C*(0) = 2 mm^3^ as per the experimental setup of Williams et al. (2000).
2. Knockout: same as (1) adding *I*(0) = 0 and *β* = 0 to prevent growth of the inflammatory compartment.
3. NSAID: same as (1) adding the concentration of NSAID *N* = 3 mg to the model, with inhibition efficacy *µ*_*nsaid*_ = 0.7 day^−1^mg^−1^, and replacing equation (2) with (7) below

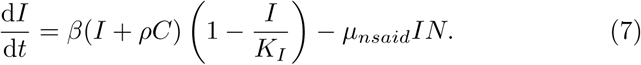

The NSAID was administered to the mice through their chow (1, 500 mg/kg celecoxib-containing chow, Williams et al. (2000)). We estimated the mouse level to be about 3 mg and assume it to be constant.

Blocking the action of COX-2 limits the ability of the carrying capacity to grow (due to VEGF suppression) and thus slows tumour growth, both experimentally and in our mathematical model. Experimental data and model simulations of the COX-2 knockout experiment are shown in Figure 4[a] and the NSAID COX-2 inhibitor treament are shown in Figure 4[b].

**Figure 4:**
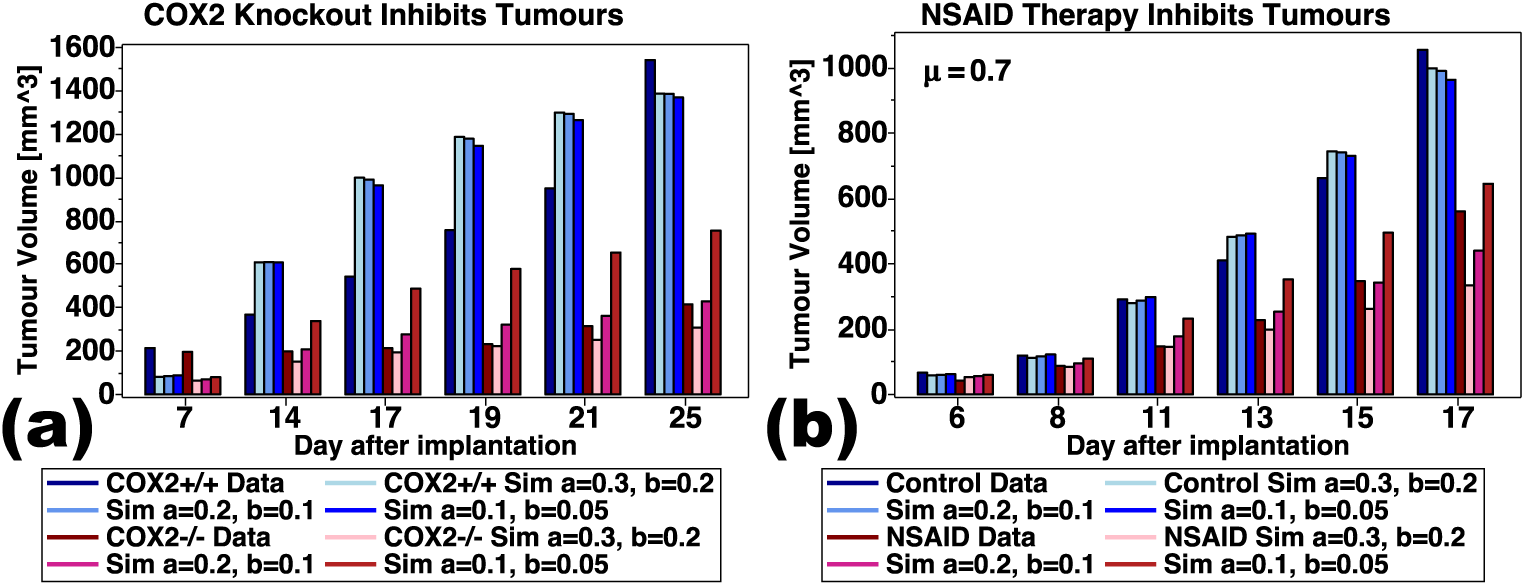
Inflammation promotes LLC tumour growth. Experimental data and model simulations of (a) the COX-2 knockout mouse experiment (Williams et al.; 2000, Figure 1a), or (b) the anti-inflammatory treatment experiment (Williams et al.; 2000, Figure 2). In (a) the LLC tumours are implanted into either COX2+/+ or COX2-/- knockout C57BL6 mice. The knockout mouse is simulated to have no inflammatory immune component *I*(0) = 0 and *β* = 0. Predicted results are for three pairs of inflammation parameters (*a, b*) which control the dependence of tumour growth on inflammatory actions: (*a, b*) = (0.3, 0.2), (0.2, 0.1), or (0.1, 0.05). In (b), anti-inflammatory treatment (Celecoxib), is dosed via 1, 500 mg/kg celecoxib-containing chow to LLC bearing mice. Simulations assume a constant blood level of celecoxib *N*(0) = 3 mg, with an efficacy of *µ*_*nsaid*_ = 0.7 day^−1^mg^−1^ acting to inhibit immune growth in equation (7). Again, model predictions compare three pairs of parameters (*a, b*) = (0.3, 0.2), (0.2, 0.1), (0.1, 0.05). Other model parameters are as listed in Table 1.

The experimental data clearly demonstrates that the LLC tumour growth is dependent on inflammatory actions, as the growth rate is inhibited by blocking COX-2. Model predictions mimic this trend, as blocking immune action (Knockout) or inhibiting immune growth (NSAID) both result in reduced tumour growth. Model predictions do not match the experimental data exactly as the basic growth parameters, Table 1, were determined from the Benzekry et al. (2017) dataset. Visual inspection suggests that the NSAID model predictions using *a* = 0.2 and *b* = 0.1 are a close approximation to the experimental data.

Other cancers may be more or less dependent on inflammation, and parameters (*a, b*) provide a way to tune the model to reflect inflammation dependence. When (*a, b*) = (0.1, 0.05), the simulated anti-inflammatory treatments are less effective, and when (*a, b*) = (0.3, 0.2), the treatment is more effective.

### 3.4 Chemotherapy and NSAID Combination Therapy

NSAID use with chemotherapy has been explored as a way to increase efficacy of treatment, Roxburgh and McMillan (2014) and others. Specifically, the combination of an NSAID (Celecoxib COX-2 inhibitor) and a chemotherapy agent (Tegafur/Gimeracil/Oteracil potassium) was recently considered by Meng et al. (2014). Here we use our model to explore the consequences of this combination treatment, and later extend it to treat the entire host.

The experiment of Meng et al. (2014) used BALB/c nude mice with a gastric cancer cell line SGC-7901. Two million cells were injected subcutaneously and after the tumour reached 5 mm in largest diameter, the mice were treated with either the NSAID, the chemotherapeutic, or the combination. Presented data showed that tumour volume decreased 30.8% with the NSAID, 50.1% with the chemotherapy, and 78.8% with the combination, on day 21 of the treatment regime, Meng et al. (2014). The treatment regime was 5 consecutive days a week for 3 weeks. The NSAID dose was 50 mg/kg, which in a 25g mouse would be about 1.25mg. The chemotherapeutic dose was 10 mg/kg, or 0.25mg per mouse.

To model these experiments we extend our single tumour model again to include effects of systemic chemotherapy (*M*(*t*)), in addition to the NSAID (*N*(*t*)). The extended model is

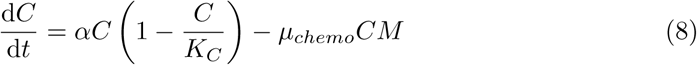

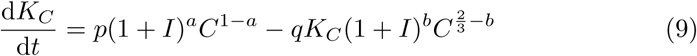

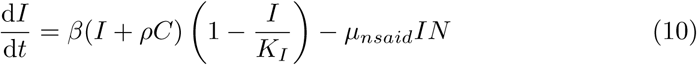

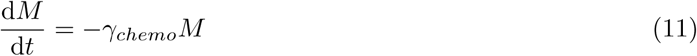

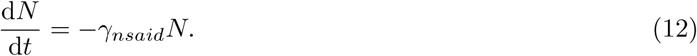

where *M* and *N* are the drug amounts in mg. The efficacy (*µ* day^−1^mg^−1^) and clearance rate (*γ* day^−1^) of the treatments are denoted as *µ*_*chemo*_ and *γ*_*chemo*_ for chemotherapy, and *µ*_*nsaid*_ and *γ*_*nsaid*_ for the NSAID. In our simulations, we choose these parameters to approximately match the single treatment inhibition results from Meng et al. (2014). Resulting parameter choices are *µ*_*chemo*_ = 0.75 day^−1^mg^−1^, *γ*_*chemo*_ = 0.48 day^−1^, *µ*_*nsaid*_ = 0.7 day^−1^mg^−1^, and *γ*_*nsaid*_ = 0.7 day^−1^, along with the parameter values of Table 1. Drug treatments are prescribed by reassigning initial conditions on days 5, 6, 7, 8, 9, 12, 13, 14, 15, 16, 19, 20, 21, 22, and 23 with additional doses of 0.25 mg for the chemotherapeutic and 1.25 mg for the NSAID. Final tumour measurements are obtained on day 25, at the end of the three week regime.

Model simulation results are shown in Figure 5. With NSAID treatment, the tumour and carrying capacity continue to grow, and as they get larger, they become stronger inducers of the immune response, which allows inflammation to grow during treatment breaks. With chemotherapy treatment, the inflammation is left unchecked, which works to promote carrying capacity growth and has the effect of increasing tumour growth rate. When both treatments are combined, the effects are almost additive in simulation. Inflammation is kept low and the tumour growth rate is reduced. Interestingly, our simulation does not predict completely additive treatment effects, in that tumour inhibition is about 30% with NSAID treatment, 50% with chemotherapy, and only 71% in combination, compared to a combined result of 79% in the experimental data. A potential explanation for this is that our model was parameterized using the LLC–C57BL6 Benzekry et al. (2017) dataset which suggested a maximal carrying capacity of 1500 mm^3^, and the experimental data of Meng et al. (2014) is reported over 2200 mm^3^. Thus, our simulations run into the strongly limiting effects of the maximal carrying capacity, which influence the third week of treatment.

**Figure 5:**
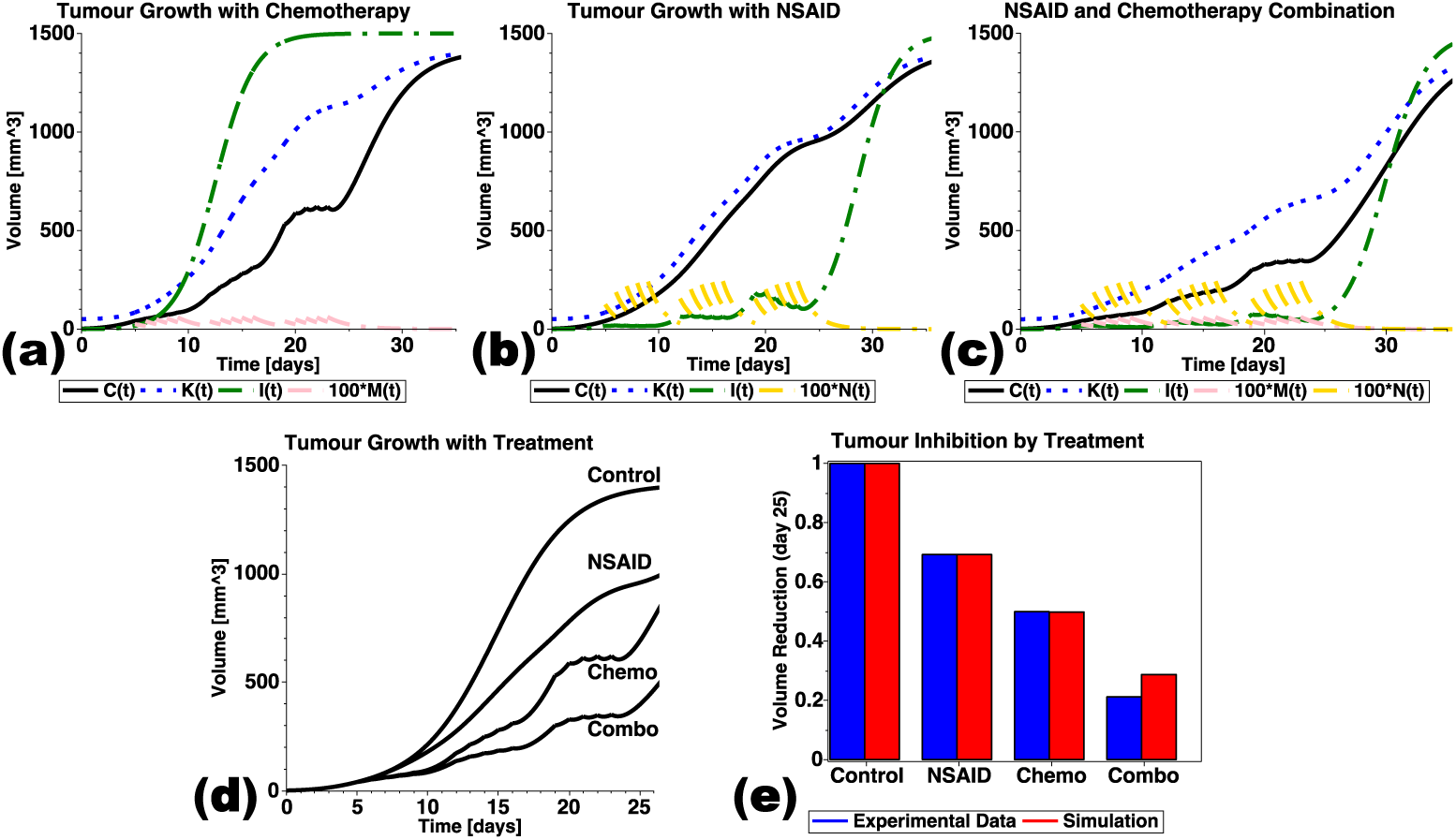
Anti-inflammatory and chemotherapy treatment for tumour control. Model simulations (equations (8)–(12)) of (a) chemotherapy (with *µ*_*chemo*_ = 0.75 day^−1^mg^−1^ and *γ*_*chemo*_ = 0.48 day^−1^), (b) NSAID therapy (with *µ*_*nsaid*_ = 0.7 day^−1^mg^−1^ and *γ*_*nsaid*_ = 0.7 day^−1^), and (c) the combination therapy demonstrate how each treament affects tumour volume, carrying capacity, and inflammatory response curves. (d) A comparison of model predicted tumour growth curves for control, NSAID, chemotherapy, and the combination treatment. (e) Comparison of predicted tumour inhibition on day 25 to the experimental data of Meng et al. (2014). If not listed, model simulations use parameter values from Table 1.

### 3.5 Treating the Whole: Chemotherapy, NSAID, and Anti-Angiogenics

The previous simulations of NSAID and chemotherapy treatment suggest a synergy exists when targeting both the tumour and host. We explore this further by adding an anti-angiogenic treatment that targets host endothelial cells and inhibits tumour carrying capacity. We extend the above model to include anti-angiogenic treatment in combination with an NSAID and chemotherapeutic. Thus, the model extended to all three treatment modalities consists of equations (8), (10)–(12) and

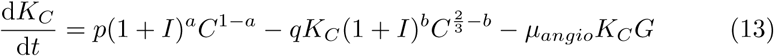

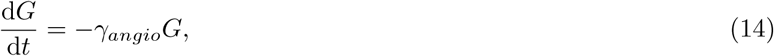

where *G* is the anti-angiogenic drug amount in mg, *µ*_*angio*_ is the efficacy, and *γ*_*angio*_ is the clearance rate.

Parameter values *µ*_*angio*_ and *γ*_*angio*_ are chosen to match model predictions to experimental data from Hahnfeldt et al. (1999), based on a dose of angiostatin at 20 mg/kg (0.5 mg per 25 g mouse) maintaining approximate tumour dormancy in the LLC–C57BL6 mouse model. The values found are *µ*_*angio*_ = 0.5 day^−1^mg^−1^ and *γ*_*angio*_ = 0.2 day^−1^.

Model predictions of treatments and combinations of treatments assume a regime of treating for 5 days on, 2 days off, for 3 consecutive weeks. Results are summarized in Figure 6. The effect of single treatments are show in Figure 6[a], and combination treatments are shown in Figure 6[b]. Insets show the tumour inhibition on day 25. Interestingly, the model suggests that targeting the inflammatory response and vascular endothelial cells is a more effective treatment strategy than either one combined with a chemotherapeutic. This finding reinforces the idea of targeting the healthy host (non-mutated) cells to control cancer. Unfortunately, in practice, anti-angiogenics have not performed as hoped due to acquired resistance and variable drug efficacy, Abdalla et al. (2018).

**Figure 6:**
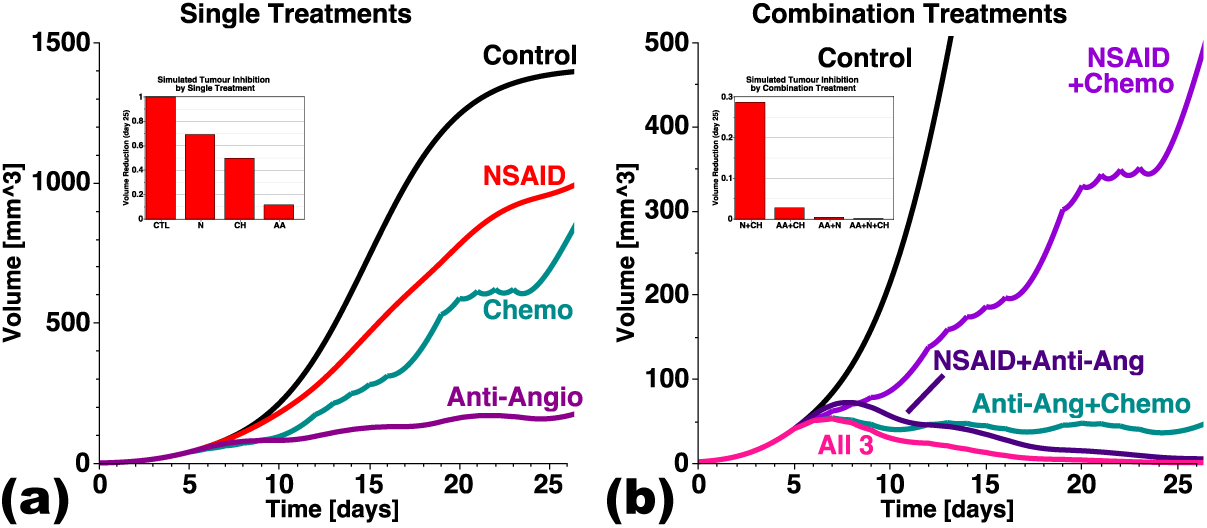
Model predictions of (a) individual treatments (chemo, NSAID, and anti-antiogenic therapies) and (b) all possible combinations. Improved tumour control is obtained by targeting both the cancer (chemotherapy) and host (anti-angiogenic and anti-inflammatory). Model simulations of equations (8), (10)–(12), (13)–(14) with parameter values listed in Table 1 and 2.

### 3.6 Modeling Simultaneous Tumour Growth

We now turn our focus to the simultaneous tumour growth experiments of Benzekry et al. (2017), where two equal LLC implants lead to one large and one small tumour, consistently. To test for biological circumstances that could lead to the growth rate separation, we fit our two-tumour model, equations (4)–(6), to the experimental data by selectively perturbing parameters associated with biological mechanisms. As baseline, we assume the control parameter values listed in Table 1. Combinations of parameters are then varied in this secondary fitting process, with a best fit determined by the minimum sum of the RMSE of each curve (i.e. minimize RMSE_big_ + RMSE_small_). The parameters/mechanisms explored assuming site 1 is the large tumour and site 2 is the small tumour are:

**Table 2:**
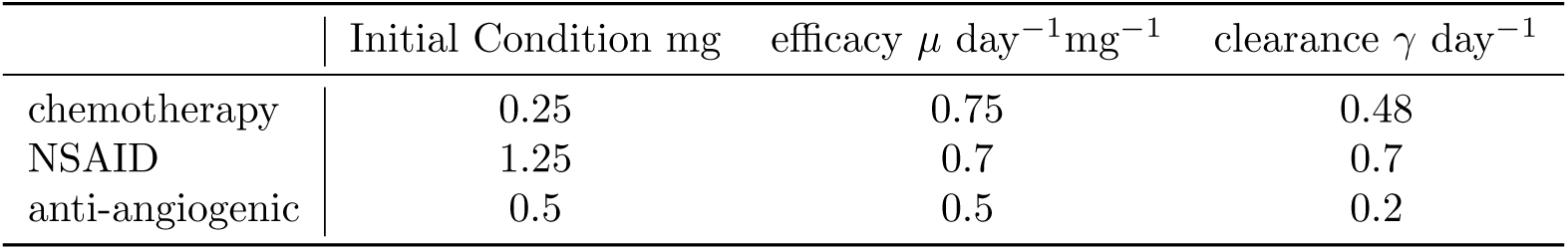
Treatment-related parameter values and initial conditions for drug dosage, efficacy, and clearance rate.

1. differences in initial tumour volume due to variations in viable injected cells (*C*_2_(0) < *C*_1_(0) with *C*_1_(0) = 10^6^ cells).
2. differences in initial immune presence (*I*_1_(0) > *I*_2_(0) = 0). Note that *I*(0) = 0 was assumed in the one tumour model.
3. differences in immune growth rate (*β*_2_ < *β*_1_). The myriad cytokines and activated immune cells may be different between the two sites causing a discreptancy in growth rates.
4. differences in immune recruitment rate (*ρ*_2_ < *ρ*_1_). The cytokine and chemokine profile may differ between the two sites and result in a discreptancy in recruitment from blood.
5. differences in initial cancer carrying capacities (*K*_*C*1_(0) > *K*_*C*2_(0)) representing, for example, variations in initial microenvironmental blood vessel density.

The results of single and combination parameter fits are summarized in Table 3. Single parameters require biologically unfeasible differences between the two sites to explain the growth separation. For example, by itself, the initial condition can fit the data, but it requires the second injection to contain only 5% viable cells compared to the first injection, which is not feasible due to the experimental set-up and replication.

**Table 3:** Results of fitting the two-tumour model, equations (4)–(6), to the experimental data of Benzekry et al. (2017), restricting only test variables to be perturbed from the control value listed in Table 1. Individual and combinations of parameters are tested.

| Test variable set-up | Result of fitting | RMSE |
| --- | --- | --- |
| $C_2(0) = \gamma C_1(0), 0 \leq \gamma \leq 1$ | $C_2(0) = 0.053 C_1(0)$ | 226.4 |
| $I_1(0) > 0, I_2(0) = 0$ | $I_1(0) = 103, I_2(0) = 0$ | 138.3 |
| $\beta_2 = \gamma \beta_1, 0 \leq \gamma \leq 1$ | $\beta_2 = 0.23 \beta_1$ | 193.4 |
| $\rho_2 = \gamma \rho_1, 0 \leq \gamma \leq 1$ | $\rho_2 = 0.33 \rho_1$ | 189.6 |
| $K_{C1}(0) > K_{C2}(0)$ | $K_{C1}(0) = 204.3 \text{ mm}^3,$<br>$K_{C2}(0) = 2.1 \text{ mm}^3$ | 146.7 |
| vary $C_2(0), I_1(0), \beta_2$, and $\rho_2$ | $C_2(0) = 0.3 \text{ mm}^3, I_1(0) = 15 \text{ mm}^3$<br>$\beta_2 = 0.9 \beta_1, \rho_2 = 0.9 \rho_1$ | 104.0 |
| vary $C_2(0), I_1(0), \beta_2, \rho_2,$<br>$K_{C1}(0)$, and $K_{C2}(0)$ | $C_2(0) = 0.5 \text{ mm}^3, I_1(0) = 1 \text{ mm}^3$<br>$\beta_2 = \beta_1, \rho_2 = 0.1 \rho_1$<br>$K_{C1}(0) = 146 \text{ mm}^3, K_{C2}(0) = 53 \text{ mm}^3$ | 99.5 |
| pre-inflamed site:<br>vary $I_1(0), I_2(0),$<br>$K_{C1}(0), K_{C2}(0)$ | $I_1(0) = 35 \text{ mm}^3, I_2(0) = 0 \text{ mm}^3,$<br>$K_{C1}(0) = 34 \text{ mm}^3, K_{C2}(0) = 18 \text{ mm}^3$ | 104.9 |

Parameter combinations provide more biologically realistic explanations as, generally, they result in smaller differences between the two sites, Figure 7 and Table 3. The best fit, Figure 7[c], allows the following parameters to differ between sites *i* = 1, 2: *C*_*i*_(0), *I*_*i*_(0), *K*_*Ci*_(0), *β*_*i*_, and *ρ*_*i*_ (RMSE=99.5). However, some of the optimal differences can be dismissed as not feasible (*C*_2_(0) = 0.5*C*_1_(0)) or as not required (*β*_2_ = *β*_1_). The fit shown in Figure 7[d] thus assumes that *C*_2_(0) = *C*_1_(0), *β*_2_ = *β*_1_ and *ρ*_2_ = *ρ*_1_, leaving only parameters *I*_*i*_(0) and *K*_*Ci*_(0) to be fit. Under these conditions, which we denote “pre-inflamed”, the resulting best fit requires *I*_1_(0) = 35 mm^3^, *I*_2_(0) = 0 mm^3^, *K*_1_(0) = 34 mm^3^, and *K*_2_(0) = 18 mm^3^ (RMSE = 104.9). That is, one site has an intial immune presence and thus a slightly larger initial cancer carrying capacity compared to the other.

**Figure 7:**
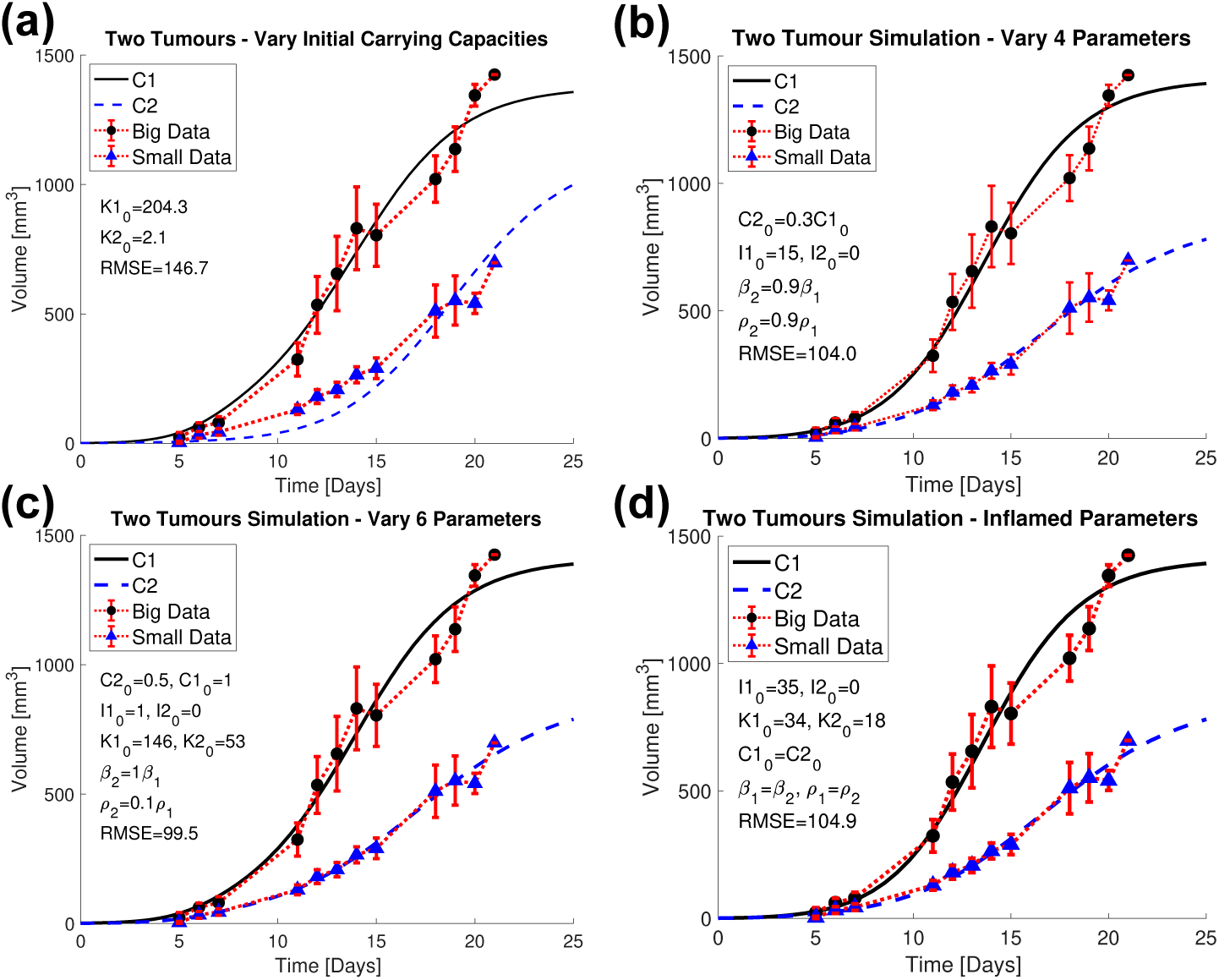
Parameter combination fits of the two-tumour model to simultaneous tumour growth data of Benzekry et al. (2017). All parameter values are those listed in Table 1, except the ones varied in the following test conditions: (a) Initial cancer carrying capacity *K*_*Ci*_(0), *i* = 1, 2, (b) Four parameter combination - *C*_*i*_(0), *β*_*i*_, *ρ*_*i*_, and *I*_*i*_(0), *i* = 1, 2, (c) Six parameter combination - the same parameters in (b) adding *K*_*Ci*_(0), *i* = 1, 2, and (d) pre-inflamed parameters *I*_*i*_(0) and *K*_*Ci*_(0), *i* = 1, 2. Parameters for best fits are listed in Table 3.

To test the concept of a pre-inflamed site, we go back to the two mouse datasets from Benzekry et al. (2017) that were initially excluded from our analysis since the mice grew tumours of equal size. One mouse grew two fast-growing tumours while the other grew two slow-growing tumours. If we assume both sites were inflamed in the two-large tumours mouse, then we should assign the pre-inflamed conditions shared between both sites: 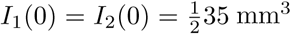 and 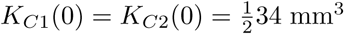. Figure 8[a] shows that model predictions are comparable to the data. Doing the same for the mouse with two slow-growing tumours, the not-inflamed parameters are shared between sites: *I*_1_(0) = *I*_2_(0) = 0 mm^3^ and 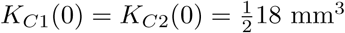. The model predicted simultaneous growth of two slow-growing tumours, comparable with the average of the small tumours, but not as slow-growing as in this single mouse.

**Figure 8:**
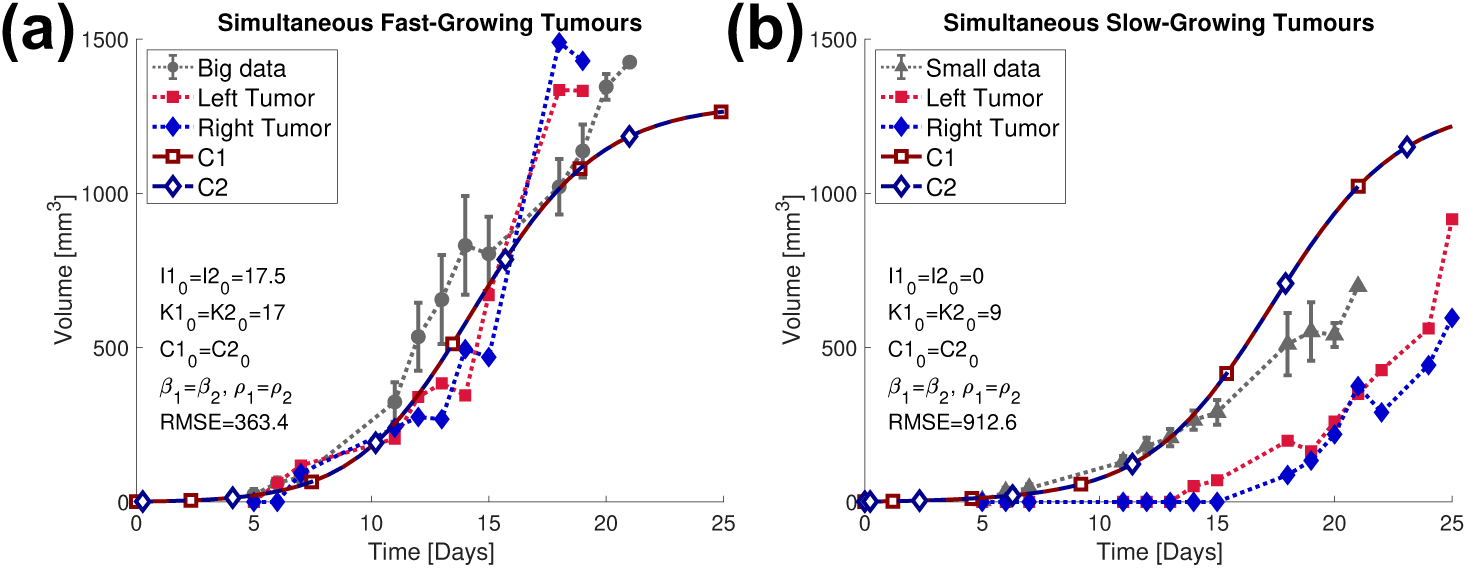
Model predictions of the two mice that grew simultaneous tumours of similar size assuming the pre-inflamed parameter values from Table 3. All parameter values are those listed in Table 1, except in (a) fast-growing tumours are assumed to share the pre-inflamed state at both sites: 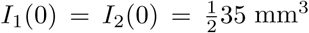 and 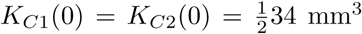 while in (b) slow-growing tumours are assumed to share the not-inflamed state at both sites: *I*_1_(0) = *I*_2_(0) = 0 mm^3^ and 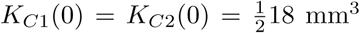.

### 3.7 Delayed Secondary Implants as a Model of Metastasis

So far we have explored the competition that occurs between tumour sites when the implants are simultaneous. Bruzzo et al. (2010) delayed the second implant for six days and altered the number of cells in the secondary injection. They saw that the second injection can be suppressed when small enough, see (Bruzzo et al.; 2010, Figure 4). Their study used a non-immunogenic cancer (LB T-lymphoid leukemia-lymphoma) in the BALB/c mouse. Experimental setup involved mice receiving 10^6^ cells on day 0, and a second injection on day 6 in the opposing flank. Controls received only the second injection on day 6. With a second injection of 10^5^ cells, no growth was observed, with 10^6^ cells, inhibited growth compared to control was observed, and with 10^7^ cells, the growth was similar to control.

We replicate this experiment with our two-tumour model by changing initial conditions to match the setup. That is, *C*_1_(0) = 1, *C*_2_(0) = 0 mm^3^, and then the system is restarted with *C*_2_(6) = 0.1, 1, or 10 mm^3^. The data of Bruzzo et al. (2010) shows inhibitory mechanisms of concommitant resistance that cannot be explained by our model of unbalanced inflammation. Note that we do not yet have data to inform the significance of inflammation in LB–BALB/c tumours. Further, Bruzzo et al. (2010) state that no host cells were detected within their secondary tumours. We thus conclude that while inflammation may contribute to systemic growth-rate interference between multiple tumour sites, alone, it cannot explain all observations of concomitant resistance.

### 3.8 The Effect of Surgery on Simultaneous Tumours

To explore the effect of surgery on cancer recurrence, Predina et al. (2012) used a simultaneous tumour growth experiment with 5 × 10^5^ AKR–esophageal cancer cells in C57BL6 mice. The two-tumour bearing mice also grew one big and one small tumour. Once the larger tumour reached 500 mm^3^ in volume, it was surgically resected. Predina et al. (2012) claim that the remaining tumour’s growth rate increases post surgery.

Using their data [Figure 1B of Predina et al. (2012)], we estimate the exponential growth rate for the small tumour using all data points, only those prior to surgery, and only those post surgery, using the ExponentialFit command in Maple. We found no increase in exponential growth rate post-surgery, see Table 4. Note that the best curve fits to the data involve excluding the *t* = 0 datapoint (data not shown), and in this case, the exponential growth rates are approximately equal pre- and post-surgery.

**Table 4:**
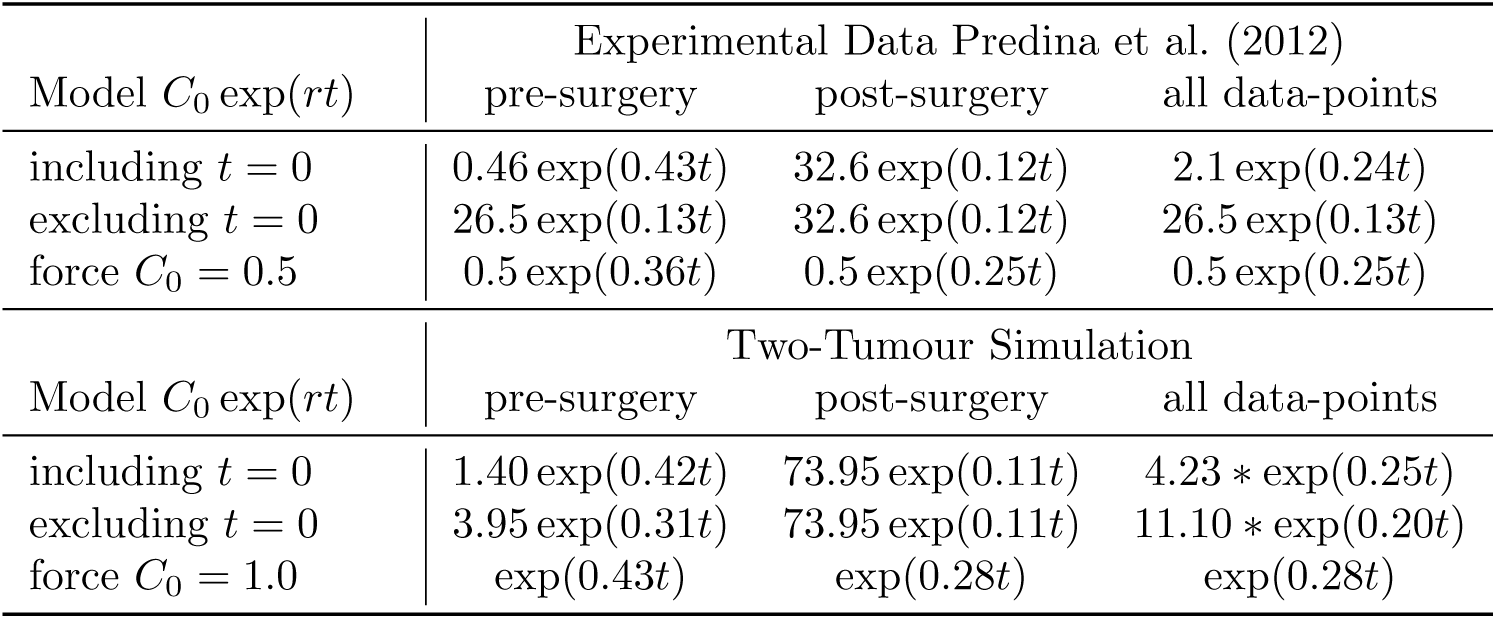
Growth rate estimation of the smaller of two tumours pre- and post-surgical removal of the larger simultaneous tumour. Experimental data of AKR implants from Predina et al. (2012) and model simulations, equations (4)–(6), with parameter values from Table 1 and the pre-inflamed condition in Table 3. Post surgery, no increase in exponential growth rate is observed, when all data points are used, when the initial *t* = 0 datapoint is excluded, or when the initial injection size is enforced.

| Model $C_0 \exp(rt)$ | Experimental Data Predina et al. (2012) | | |
| --- | --- | --- | --- |
|  | pre-surgery | post-surgery | all data-points |
| including $t = 0$ | $0.46 \exp(0.43t)$ | $32.6 \exp(0.12t)$ | $2.1 \exp(0.24t)$ |
| excluding $t = 0$ | $26.5 \exp(0.13t)$ | $32.6 \exp(0.12t)$ | $26.5 \exp(0.13t)$ |
| force $C_0 = 0.5$ | $0.5 \exp(0.36t)$ | $0.5 \exp(0.25t)$ | $0.5 \exp(0.25t)$ |
| Model $C_0 \exp(rt)$ | Two-Tumour Simulation | | |
|  | pre-surgery | post-surgery | all data-points |
| including $t = 0$ | $1.40 \exp(0.42t)$ | $73.95 \exp(0.11t)$ | $4.23 * \exp(0.25t)$ |
| excluding $t = 0$ | $3.95 \exp(0.31t)$ | $73.95 \exp(0.11t)$ | $11.10 * \exp(0.20t)$ |
| force $C_0 = 1.0$ | $\exp(0.43t)$ | $\exp(0.28t)$ | $\exp(0.28t)$ |

Nevertheless, we mimic this experiment using our pre-inflamed parameter set from Table 3 and base model parameters from Table 1, in the two-tumour model, equations (4)–(6). Surgery is simulated on day 12 to be 100% effective at removing the larger tumour, *C*_1_(*t*), but only 99% effective at clearing the cancer capacity and immune presence at site 1. Timepoints used to estimate the exponential growth rate are *T* = [0, 5, 8, 10, 12, 16, 18, 22, 24, 26], giving 5 datapoints pre- and post-surgery, as in the experimental setup. Similar to the data, we find no increase in exponential growth rate post surgery.

### 3.9 Inflammatory Effects in Cancer Recurrence

Inflammation can promote cancer metastases, increasing the risk of recurrence. For example, breast cancer death risk curves have a single peak for untreated patients but have two peaks for treated patients, Galmarini et al. (2014). It has been hypothesized that the early peak corresponds to the removal of con-commitant resistance mechanisms linked to the primary tumour and to the activation of dormant metastases by surgically-induced inflammation, while the second peak corresponds to recurrence due to a stochastic switch out of dormancy, Galmarini et al. (2014). Indeed, Krall et al. (2018) showed that surgical wounding at a distant site promotes tumour growth and that the use of NSAID Meloxicam can counteract this stimulation. They propose that NSAID use could reduce the early metastatic recurrence peak in breast cancer.

We use our two-tumour model, equations (4)–(6), to explore the effects of surgery and NSAID use on a secondary tumour implanted 10 days after the primary. Each tumour is simulated to grow from 1 cell (*C*_*i*_(*t*^∗^) = 10^−6^ mm^3^) and is assumed to have an initial carrying capacity of ten times the initial cancer size, *K*_*Ci*_(*t*^∗^) = 10*C*_*i*_(*t*^∗^) where *t*^∗^ = 0 for the primary at site *i* = 1 and *t*^∗^ = 10 for the metastasis at site *i* = 2. Other parameter values are those listed in Table 1. Surgery occurs on day 50 at site *i* = 1 with clearance efficacy of 100% for the primary tumour volume and 95% for immune presence and cancer carrying capacity.

Model prediction of the primary and metastasis tumour volumes with surgerical removal of the primary is shown in Figure 10[a]. Surgery frees up inflammatory cells from the primary site which then relocate to the metastasis, enhancing it’s maximal size. The relative increase in metastatic volume post surgery is shown in Figure 10[b]. Relative increase is calculated by the formula 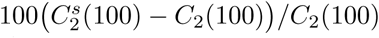 where *C*_2_(100) is the metastasis volume with the primary present at *t* = 100, and 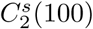 is the metastasis volume post surgery at *t* = 100 days. As a function of surgical date, relative increase has a non-monotonic shape, suggesting there is an optimal surgical date (around day 30) where metastatic enhancement post-surgery is minimized. This effect changes, however, as surgical efficacy is reduced, in which case, earlier surgeries result in smaller metastases because of primary tumour recurrence (95% efficacy).

**Figure 9:**
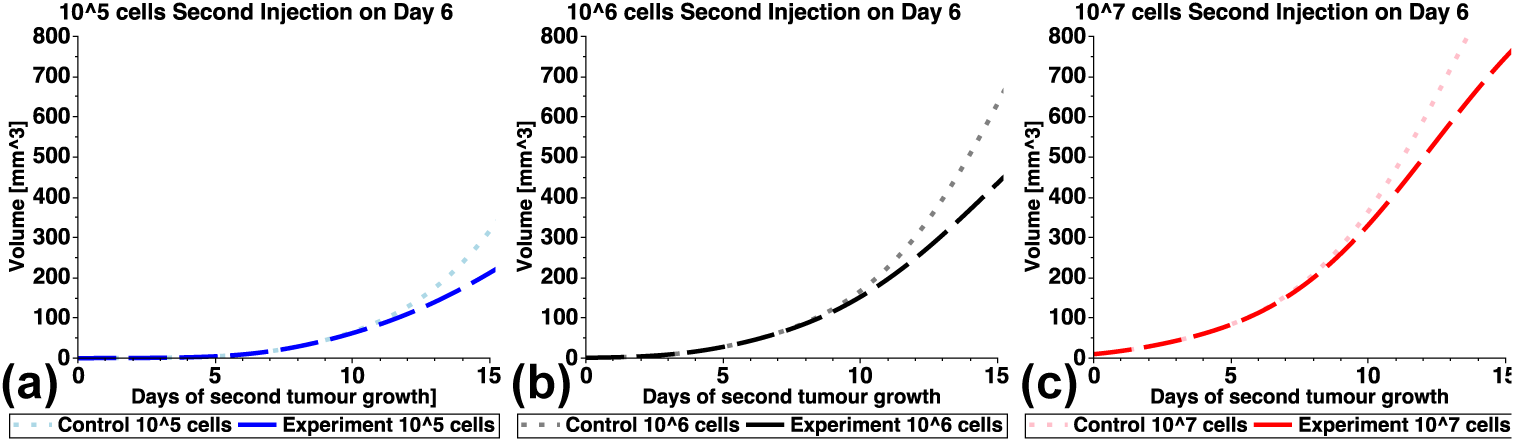
The effect of delaying the second tumour injection. Model simulation is started with one tumour *C*_1_(0) = 1 mm^3^ on day 0 and a second tumour is started on day 6 with either (a) *C*_2_(6) = 0.1, (b) *C*_2_(6) = 1, or (c) *C*_2_(6) = 10 mm^3^. Predictions show all secondary tumours experience delayed growth compared to the control, which is a single tumour of the same size started on day 6. Model simulations use parameter values listed in Table 1.

**Figure 10:**
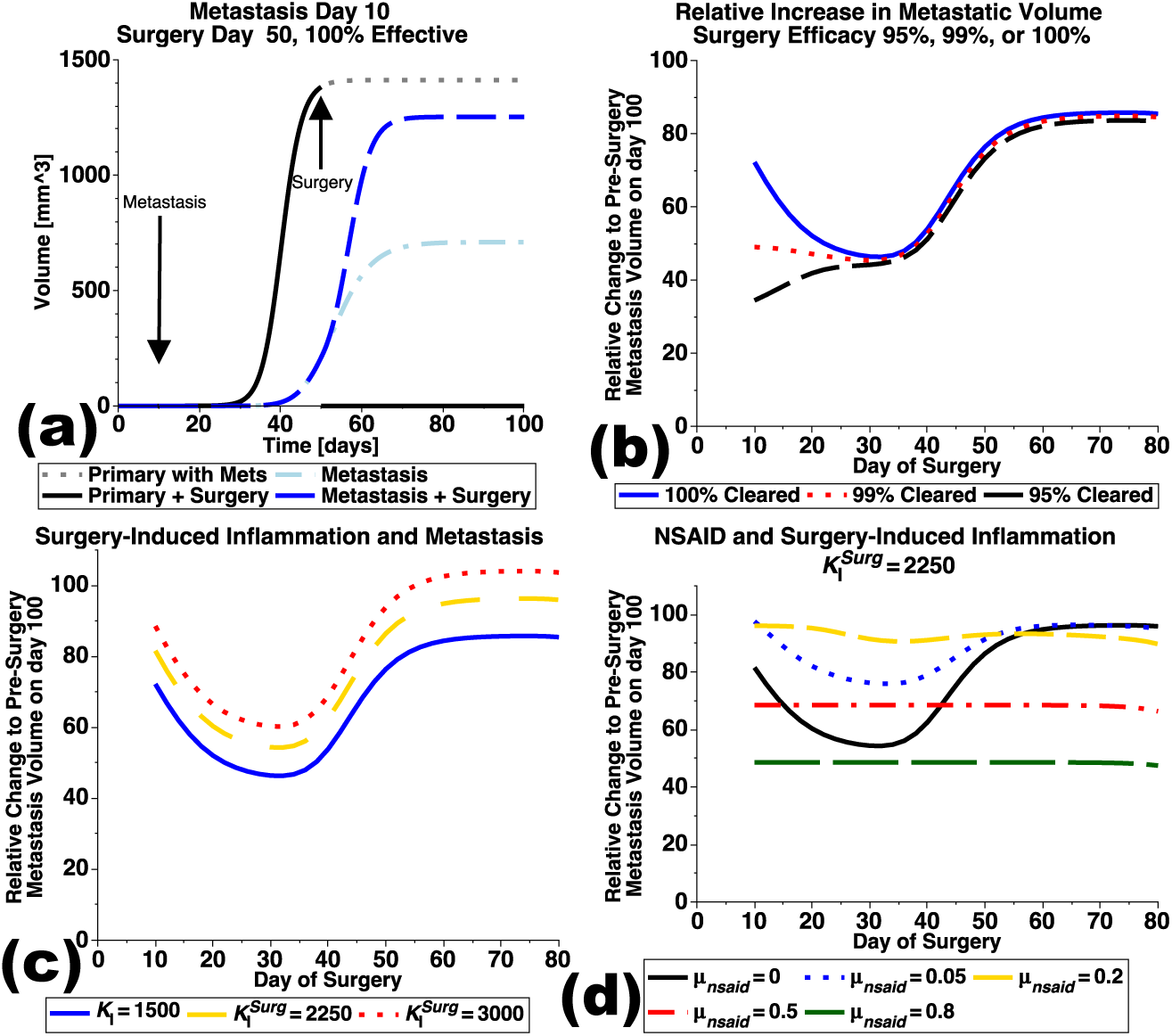
Model predictions of metastasis growth post surgical removal of the primary. (a) Simulation of primary growth from *C*_1_(0) = 10^−6^ mm^3^, metastasis started on day 10, *C*_2_(10) = 10^−6^ mm^3^, with surgery on day 50 removing 100% of the primary *C*_1_, but only 95% of *K*_*C*1_ and *I*_1_. Post surgery, immune cells are relocated from site 1 back to the systemic pool, where they can move to site 2 and enhance the maximal size of the metastasis. (b) Changing surgery date alters the metastatic enhancement, with a minimum occuring near day 30 for 100% primary clearance. Less effective surgeries reduce metastatic enhancement when performed early on, but result in similar enhancement when performed late. (c) Surgical stimulation of systemic inflammation increases the metastatic enhancement and maintains the nonmonotonic response to surgical date. Simulations performed by increasing *K*_*I*_ by a factor of 1.5 or 2 after surgery. (d) Anti-inflammatory treatment with surgical enhancement of systemic inflammation. Strong NSAID efficacy flattens the non-monotonic curve and reduces metastatic enhancement post surgery. Metastatic enhancement post surgery computed via 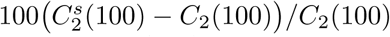, where *C*_2_(100) is metastasis volume with primary and 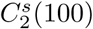 is metastasis volume post surgery, on day *t* = 100. Unless otherwise stated, parameter values are from Table 1.

Next, we allow the act of surgery to stimulate an enhanced inflammatory response. Surgical efficacy is fixed at 100%, and surgery increases *K*_*I*_ by a factor of 1.5 or 2. Figure 10[c] shows that increasing *K*_*I*_ post surgery enhances the metastasis, but maintains the non-monotonic response to surgery date. Finally, we add an anti-inflammatory treatment, by adding −*µ*_*nsaid*_*I*_*i*_(*t*)*N*(*t*), *i* = 1, 2 to the right hand side of equation (6), similar to equation (10), with *N*(0) = 1.25 mg as used above for Celecoxib. For simplicity, we assume NSAID dosing is such that a constant blood level is maintained, *N*′ = 0. Interestingly, the anti-inflammatory treatment can remove the effect of surgical date on metastasis enhancement if the efficacy, *µ*_*nsaid*_, is high enough, Figure 10[d].

## 4 Discussion

We presented a mathematical model that describes stimulation of tumour growth due to systemic inflammation. Significant model parameters were estimated by fitting to experimental data of Benzekry et al. (2017), others were fixed pre-fitting. Model parameters *a* and *b* control the weight of pro- and anti-tumour effects the immune cells have on tumour carrying capacity. Using additional data from Williams et al. (2000), we validated the choice of *a* = 0.2 and *b* = 0.1 for the dependence of LLC–C57BL6 tumour growth on inflammation. Other tumour models will require different values for *a* and *b*.

Using our model for tumour, tumour carrying capacity, and immune components, we incorporated chemotherapy, anti-inflammatory, and anti-angiogenic treatments. Parameters for drug clearance and efficacy were estimated from experimental data. Model predictions suggest that targeting the host immune response and endothelial (angiogenic) cells results in the best 2-drug combination result. As is expected, using all 3 treatments gives the best tumour control. Next, we extended the model to explore the role of inflammation in simultaneous tumour growth with growth-rate separation. Our hypothesis was that inflammation may contribute to the growth rate disparities between tumour sites by preferentially accummulating at one site over the other. Thus adding another potential mechanism for concommitant resistance in tumours dependent on inflammatory processes. Parameter values for the linked two-tumour model were estimated from experimental data of Benzekry et al. (2017), and the most plausible explanation from model fitting was determined to be that the larger tumour site was pre-inflamed compared to the other.

Our two-tumour model is a biologically-motivated mathematical description requiring relatively few model parameters that is capable of describing the growth rate separation observed in the experimental data. This modelling work suggests that an injury to the site prior to injection may explain the growth rate separation observed. Notably, since we developed the mathematical model in the presence of tumour-promoting inflammation, we did not need to introduce any inhibitory mechanism to explain the growth suppression of one of the tumours. Our suggestion of a previously inflamed site seems plausible given that the experimental animals are not isolated and that they are generally quite active. These two factors increase the likelihood of them acquiring a bruise or other small injury in the general area of injection, pre-experiment. We note that the experimental mice were male, which are more active and aggressive, in general, than female mice.

Concomitant resistance was previously explored mathematically by Benzekry et al. (2017) to describe the growth rate separation observed in this data. Due to the time-lags (and tumour sizes) generally associated with CR, Chiarella et al. (2012); O’Reilly et al. (1994), however, we suggest an alternate mechanism to explain this simultaneous injection data. Using our model to replicate the delayed secondary injection experiments of Bruzzo et al. (2010), however, demonstrates that while inflammation may contribute to the growth rate interference, it cannot explain all observations of concommitant resistance.

Concomitant resistance is typically attributed to the observation of accelerated metastatic growth upon resection of the primary tumour, Chiarella et al. (2012). We add that accelerated metastatic growth may also be attributed to an increase in inflammatory cells locating to metastatic sites upon surgical injury and removal of the primary tumour. Such a biological mechanism would act in concert with the previously established CR mechanisms of immune cytotoxicity, nutrient hoarding, and angiogenesis or proliferation inhibitors.

To explore the effects of surgery on metastatic growth, we use our two-tumour model and compare simulations to experimental data of Predina et al. (2012). Exponential growth rates were estimated from both the data and model simulations but no increase in growth rate was observed in either dataset.

However, post-surgery metastatic enhancement of tumour volume was observed. Interestingly, a non-monotonic relationship was found between metastatic enhancement and surgery date, with later surgeries resulting in maximal enhancement, and a minimal enhancement occuring near day 30 when surgery was assumed 100% effective. When surgery was assumed to increase systemic inflammation, *K*_*I*_, the model predicts an increase in the metastatic enhancement due to removal of the primary. Addition of anti-inflammatory NSAID treatment, however, was predicted to flatten the metastatic enhancement curve due to surgery, and reduce enhancement for strong drug efficacy.

As more data and model predictions are generated demonstrating the influence of inflammation on cancer growth and progression, standards of care may move to include anti-inflammatory agents. Such combination therapies would be wholistic in the sense that they target both the cancer (chemotherapy) and the host’s responses to cancer presence (anti-angiogenic and anti-inflammatory therapies). In particular, as suggested by Roxburgh and McMillan (2014), blocking the inflammatory response by treating both the tumour and the host, may have significant implications in patient quality of life, with reduced pain and reduced weight loss, leading to improved survival. The development and analysis of predictive mathematical models can help to further our understanding of the destructive effects of a systemic inflammatory response in cancer and to inform optimal therapeutic approaches.

## Acknowledgements

The authors would like to thank Dr. S. Benzekry, C. Lamont, Dr. L. Hlatky, and Dr. P. Hahnfeldt for their helpful discussions and support from the Centre of Cancer Systems Biology (http://cancer-systems-biology.org). Funding support for KPW was provided by an NSERC Discovery Grant.

